# Development of a genome scale metabolic model for the lager hybrid yeast *S. pastorianus* to understand evolution of metabolic pathways in industrial settings

**DOI:** 10.1101/2023.10.25.564032

**Authors:** Soukaina Timouma, Laura Natalia Balarezo-Cisneros, Jean-Marc Schwartz, Daniela Delneri

## Abstract

*In silico* tools such as genome-scale metabolic models (GSMM) have shown to be powerful for metabolic engineering of microorganisms. Here, we created the iSP_1513 GSMM for the aneuploid hybrid *S. pastorianus* CBS1513 to allow top-down computational approaches to predict the evolution of metabolic pathways and to aid strain optimisation and media engineering in production processes. The iSP_1513 comprises 4062 reactions, 1808 alleles and 2747 metabolites, and takes into account the functional redundancy in the gene-protein-reaction rule caused by the presence of orthologous genes. Moreover, a universal algorithm to constrain GSMM reactions using transcriptome data was developed as a python library and enabled the integration of temperature as parameter. Essentiality datasets, growth data on various carbohydrates and volatile metabolites secretion were used to validate the model. Overall, the iSP_1513 GSMM represent an important step towards understanding the metabolic capabilities, evolutionary trajectories and adaptation potential of *S. pastorianus* in different industrial settings.

## INTRODUCTION

Over the last decade, many genome-scale metabolic models of *S. cerevisiae* have been constructed (Chen *et al.*, 2022) and have become increasingly popular over as they provide a comprehensive view of the metabolic network and enable the prediction of cellular behaviour under different conditions. It successfully helped elucidate new biological processes, and design cell factories producing compounds of interest (Lopes & Rocha, 2017). *S. cerevisiae* related species and hybrids also benefit from a high biotechnological interest. In fact, natural hybrids between *Saccharomyces* species are found in many industrial situations, particularly in beer brewing and in wine making, where hybrid strains evolve due to adaptation to selective environmental conditions (Alsammar & Delneri, 2020; García-Ríos & Guillamón, 2019; Krogerus *et al.*, 2018).

*S. pastorianus* is an allopolyploid sterile hybrid of the mesophilic *S. cerevisiae* and the cold tolerant *S. eubayanus*. This hybrid species has evolved naturally in industrial settings thanks to its ability to ferment at low temperature (bottom-fermenting lager yeast) under stressful conditions (Monerawela & Bond, 2017a). *S. pastorianus* is very efficient to consume and ferment the wide range of sugars found in wort and must, in addition to producing an aroma profile interesting for brewers (Giannakou *et al.*, 2021; Mortimer, 2000). Specifically, *S. pastorianus* is known for its ability to carry out alcoholic fermentation, converting sugar into ethanol and carbon dioxide, and to produce a range of esters, which are responsible for fruity and floral flavours in beer (Briggs, 2004). Beside their industrial importance, *S. pastorianus* can be a key model for studying the evolution of hybrid genomes as this species offers a unique perspective on the genomic alterations that may occur after a recent allopolyploidisation event. Comparative genomics analysis highlighted the presence of a complex structural variation among *S. pastorianus* strains, including hybrid genes and evidence of breakpoint re-usage (Hewitt *et al.*, 2014; Monerawela & Bond, 2017b). Genomics studies on four *S. pastorianus* strains, CBS 1513, CBS 1503, CBS 1538 and WS 34/74, revealed that the low amount of genetic redundancy comes primarily by the loss of the *S. cerevisiae*-like genes (Timouma *et al.*, 2020), and during fermentation the expression of orthologues is positively correlated with gene copy number (de la Cerda Garcia-Caro *et al.*, 2022). It has also been shown that the functional redundancy, generated by the presence of orthologous parental genes, is discouraged, a scenario that fits well with the gene balance hypothesis (Timouma *et al.*, 2021).

Widening the computational modelling methods established for *S. cerevisiae* reference strain to other hybrids is key to study the evolution of hybrid genomes as well as aid media and strain engineering towards the optimisation/minimisation of desirable/undesirable flavour compounds (Bizaj *et al.*, 2012). Evolutionary biology studies are already exploiting genome scale metabolic models to predict adaptive trajectories in *Escherichia coli*, which could help understand mechanisms behind speciation and adaptation (Großkopf *et al.*, 2016; Yazdanpanah *et al.*, 2023). Predictive quantitative approaches have not been used so far with *S. pastorianus* strains due to a lack of molecular data and to the multiplicity of environmental and physiological factors that influence these yeast hybrids during the fermentation process. Indeed, while the sequence of *S. cerevisiae* genome has been available for more almost 30 years, only more recently the hybrid genomes of some *S. pastorianus* strains have been sequenced and annotated (De León-Medina *et al.*, 2016; Hewitt *et al.*, 2014; Nakao *et al.*, 2009; Okuno *et al.*, 2016; Timouma *et al.*, 2020; van den Broek *et al.*, 2015; Walther *et al.*, 2014).

Here, we present iSP_1513, the first GSMM of the industrial Group I strain *S. pastorianus carlsbergensis* (CBS 1513), which was originally isolated by the Carlsberg Laboratory in Denmark in the late 1800s and is commonly referred to as the "Carlsberg yeast". This strain is known for its ability to produce clean, crisp, and refreshing lagers with a smooth mouthfeel, and is used by many breweries around the world to produce a wide range of lager styles (Gibson & Liti, 2015). iSP_1513 model offers a platform to determine and validate optimal growth conditions for specific traits of industrial relevance, such as maltose utilisation, leucine amino acid catabolism (*i.e.,* involved in the production of aromatic flavour compound) or ethanol tolerance. Flux Balance Analysis (FBA) and Flux Variability Analysis (FVA) have been used to predict growth in different environments. The effect of high and low temperatures on metabolite production have been investigated and a python library has been developed to map transcriptome data. A preference of cellular function between parental alleles has been found, with the oxidative and non-oxidative parts of the pentose phosphate pathway primarily carried out by the *S. eubayanus*-like and *S. cerevisiae*-like genes, respectively. We found that some reactions in this pathway show a shift of parental allele usage according to the temperature. Overall, the iSP_1513 GSMM and the tools developed for this hybrid will aid the prediction of fluxes in metabolic pathways in different environments, ultimately important for strain development, media engineering and to understand adaptation processes.

## RESULTS AND DISCUSSION

### Genome scale model draft reconstruction

The Yeast Consensus Model (Yeast8) provides a highly curated and comprehensive representation of yeast metabolism (Lu *et al.*, 2019), and has been used as a template to reconstruct the draft genome scale metabolic model of *S. pastorianus*. The reactions in Yeast8 that are present/absent in *S. pastorianus* were identified based on the presence/absence of the genes that support these reactions. Among the 9,728 ORFs of *S. pastorianus*, 3,742 and 5,219 are *S. cerevisiae*-like genes and *S. eubayanus*-like genes, respectively, while 1,708 pairs of orthologous parental genes are functionally redundant (Timouma *et al.*, 2020).

Firstly, the *S. cerevisiae*-like genes of *S. pastorianus* have been mapped to the genes of Yeast8. Out of the 3,742 *S. cerevisiae*-like genes of *S. pastorianus*, 748 were identified as present in Yeast8. Secondly, to map the *S. eubayanus*-like genes of *S. pastorianus* to the Yeast8 genes, one-to-one orthologs between the *S. eubayanus*-like genes and *S. cerevisiae* S288C genes have been searched using the HybridMine software (Timouma *et al.*, 2020). As a result, out of the 5,219 *S. eubayanus*-like genes, 2,536 were found to have a one-to-one ortholog in *S. cerevisiae* S288C with more than 70% identity (2,015 shared more than 80%). After this analysis, 622 *S. eubayanus*-like genes had a 1:1 ortholog present in the Yeast8 model. In total, 946 out of the 1,160 genes of Yeast8 were found to be also present in *S. pastorianus*, of which 424 have both parental genes, 324 only have an *S. cerevisiae*-like gene associated, and 198 only have the *S. eubayanus*-like gene associated, whereas 214 genes present in Yeast 8 did not have any 1:1 orthologs in *S. pastorianus*.

The absence of this pool of genes could either be biologically genuine (*i.e.,* specific gene loss in *S. pastorianus*) or a technical artefact due an incomplete functional annotation of *S. pastorianus* genes. In fact, the functional annotation was initially performed using the gene sequences of *S. pastorianus* against the gene sequences of the parents *S. cerevisiae* and *S. eubayanus* (Timouma *et al.*, 2020). To strengthen the annotation, the protein sequences were used to search for one-to-one orthologs between the *S. pastorianus* proteins and the two parental proteomes. Following this analysis, we were able to map a further 158 genes from the pool of the 214 Yeast8 genes initially not identified in *S. pastorianus*. As expected, all the *S. cerevisiae*-like and *S. eubayanus*-like genes were also predicted as *S. cerevisiae*-like and *S. eubayanus*-like proteins, respectively. Moreover, we identified inconsistencies between the genes and proteins annotation. For example, 29 and 9 *S. cerevisiae*-like and *S. eubayanus*-like genes, respectively, had a different annotation associated to their proteins, because HybridMine attributed different isoforms at gene and protein level (Supplementary Table 1). For instance, the *S. pastorianus* ORF SPGP0D00350 was predicted to be the *S. cerevisiae*-like YPR043W (RPL12A) at the gene level, while at the protein level it was predicted to be the *S. cerevisiae*-like YJR094W-A (RPL12B). Here, these two paralogs share 100% identity at protein level, hence cannot be discriminated by the software.

In summary, out of the 1160 genes of Yeast8, 1104 were found to be also present in *S. pastorianus*, of which 681 have both parental genes, 72 only have an *S. cerevisiae*-like gene associated, and 351 only have the *S. eubayanus*-like gene associated (Figure 1). In total, 56 genes were removed as they were not present in *S. pastorianus* genomic data (Supplementary Table 2), as well as 12 reactions that are exclusively supported by these genes. Additionally, 38 *S. pastorianus* genes were removed from the model as they were not associated to any reaction (Supplementary Table 2). In total, 94 genes were removed from Yeast8.

**Figure 1:**
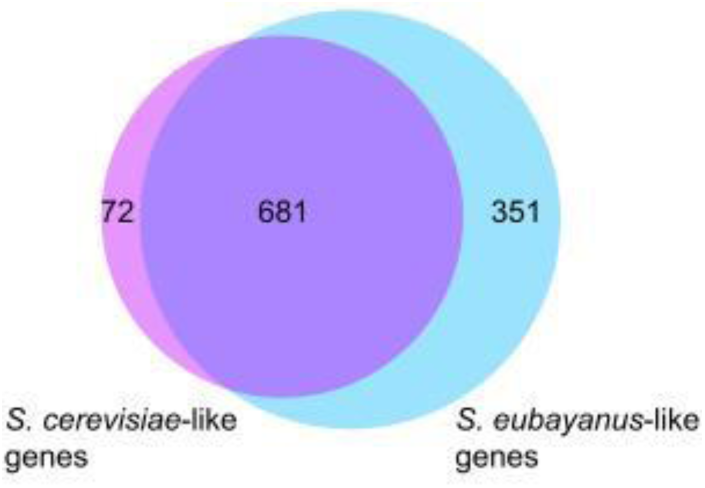
Venn diagrams showing the proportion of *S. cerevisiae*-like (purple) and *S. eubayanus*-like (light blue) genes found in the Yeast8 genome scale model. The intersection represents the proportion of genes that have both parental copies in *S. pastorianus*.

Gene enrichment was carried out on the 56 *S. cerevisiae* specific genes that were removed from Yeast8. Among them, 10 are involved in transport of sugar (glucose, D-galactose), amino-acid (L-lysine, L-Aspartate), ion (thiamine, P-type cation (Cadmium), sodium) and water (Supplementary Figure 1). In fact, as membrane proteins are under strong adaptive selection (Sojo *et al.*, 2016), it is not surprising that this class of protein is represented among the proteins specific to *S. cerevisiae* and absent in *S. pastorianus*. Moreover, it has been shown that the gene *SUL1*, encoding a sulfate permease 1, is not functional in *S. pastorianus* (Libkind *et al.*, 2011). This enzyme is indeed present in the list of genes removed (Supplementary Table 2). Finally, when at least one subunit of a protein complex was not identified as present in *S. pastorianus* genome, the hole protein complex was removed. The list of removed genes includes indeed, among others, all the genes involved in protein complex supporting mitochondrial reactions, since the mitochondria genome was not represented in the genomic data (*i.e.,* please note that the majority of these mitochondrial genes were re-integrated in the model in the follow-up manual curation, see section “Genome scale model manual curation”). This initial model, named iSP_1513v_0_, represented the first step towards drafting the *S. pastorianus* genome scale metabolic model.

### Genome scale model manual curation

Manual curation of a GSMM involves the addition, removal, and modification of reactions and associated genes and metabolites to improve the accuracy and predictive capabilities of the model. Among the reactions that were removed, 7 were re-integrated as *S. pastorianus* possesses the enzyme isoforms supporting those reactions (Supplementary Table 3). In fact, *S. pastorianus* gene SPGP0AH00790 has been predicted at the gene level as *S. eubayanus*-like *HXT5* gene while the protein was predicted to be Hxt10p. One reaction in Yeast8 involves uniquely Hxt10p while 5 reactions involve both isoforms. These 6 reactions have been re-integrated to iSP_1513 as SPGP0AH00790 can support them. *S. pastorianus* gene SPGP0EQ00100 has been predicted at the gene level as a *S. cerevisiae*-like *FDH2* (truncated protein), while the protein was predicted to be Fdh1p. Only one reaction in Yeast8 involves Fdh1p. Again, as SPGP0EQ00100 can support this reaction, this was re-integrated.

Additionally, among the initial 56 genes removed from Yeast8, the mitochondrial genes *ATP8*, *ATP6*, *COB*, *OLI1*, *COX1*, *COX2* and *COX3* were present (Supplementary Table 2), since only the nuclear genomic data of *S. pastorianus* was available. Given that the presence of functional mitochondrial DNA in *S. pastorianus* cells has been confirmed (Zhang *et al.*, 2022), the Yeast8 mitochondrial reactions, initially removed, were investigated to understand whether they could be supported by existing *S. pastorianus* enzymes. As result, four reactions were re-introduced: *i.* as *S. pastorianus* CBS1513 genome contains the *S. eubayanus*-like gene *APA2* (ID: SPGP0Y01780) that supports the ATP adenylyltransferase reaction (ID: r_0222), the reaction and gene were both added to iSP_1513; *ii.* the mitochondrial ATP Synthase “r_0226” reaction was reintroduced since *ATP6*, *ATP8* and *OLI1* genes are encoded by the mitochondrion and all the other subunits of this protein complex are present (*i.e.,* a total of 18 genes were added to the reaction GPR of which 5 have both parental genes); *iii.* the Yeast8 mitochondrial ferrocytochrome-c: oxygen oxidoreductase “r_0438” reaction was re-introduced since is supported by the mitochondrially encoded genes *COX1*, *COX2*, and *COX3* and the other subunits of this protein complex are present (*i.e.,* a total of 17 genes were added of which 5 have both parental genes; *iv.* the Yeast8 mitochondrial ubiquinol: ferricytochrome c reductase “r_0439” reaction was reintroduced as the *COB* gene is encoded by the mitochondrion and the other subunits of this protein complex are present (*i.e.,* 11 genes were added of which 5 have both parental genes).

Next, the reaction specific to *S. pastorianus*, not present in Yeast8, were investigated by mining published data and by employing KBase, a software that offers methods to construct draft genome scale models. The investigation of published data gave limited insights, but clarified a few cases. For example, unlike *S. cerevisiae*, *S. pastorianus* possesses the ability to use melibiose as carbon source, an ability inherited from the *S. eubayanus* parental sub-genome. The alpha-galactosidase (also known as melibiase) required to break the melibiose into a glucose and a galactose is encoded by the gene *MEL1*, and is not present in Yeast8. The new reaction (ID: r_4711) as well as a new gene (ID: SPGP0R03440; Name: MEL1_Seub) were added to iSP_1513.

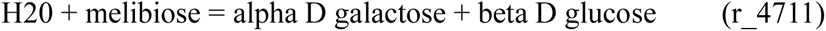

Next, we built an independent *S. pastorianus* GSMM draft using KBase. The resulting model was manually updated to take into account the functional redundancy of *S. pastorianus*. This updated model contains 1206 reactions, 1228 metabolites and 1207 genes. The *S. pastorianus* genes and reactions that are present in the KBase draft but absent from the draft iSP_1513v_0_ were identified. Because of the heterogeneity of reaction names between the iSP_1513v_0_ and the KBase models, regular expressions were used to catch key words within reaction names. As result, 1046 and 45 KBase reactions and genes, respectively, were found to be absent from iSP_1513v_0_, and therefore could be potentially added (Supplementary Table 4). Out of the 45 genes, 23 are *S. cerevisiae*-like alleles, 17 are *S. eubayanus*-like alleles and 5 *S. pastorianus* specific). The reactions supported by these genes were mapped to the iSP_1513v_0_ GSMM to identify genes/reactions specific to *S. pastorianus*. In total, 5, 26 and 3 reactions, genes and metabolites were added, respectively. The reactions/genes are detailed in Supplementary File 1.

### GPR rules update for iSP_1513

Since *S. pastorianus* carries an aneuploid and hybrid genome, genes can have *S. cerevisiae*-like alleles and *S. eubayanus*-like alleles, which results in functional redundancy. Therefore, the gene-protein-reaction (GPR) rules of Yeast8 are not applicable to *S. pastorianus* as this functional redundancy needs to be considered to enable gene deletion simulations or integration of gene expression data. Metabolic reactions can be non-enzymatic, when catalysed by small molecules, or enzymatic, when catalysed by specific proteins (Filippo *et al.*, 2021). From a structural point of view, enzymes can either be either monomeric, meaning they are composed of a single gene product, or oligomeric, meaning they are composed of multiple gene products. In the context of predicting the effects of gene deletion on metabolic fluxes, it is important to consider functional redundancy in the genome scale models, including type and composition of catalysts. This is particularly relevant in hybrid species where both parental genes are competing. GPRs are Boolean logic relationships between gene products, such as enzyme isoforms or subunits, involved in catalysing a particular reaction. The presence of functionally redundant parental alleles may impact on the nature of protein complexes established in the hybrid, where both parental alleles are competing. The protein complexes in *S. pastorianus* can be either exclusively uni-specific with subunits coming only from one parent; or exclusively chimeric, with a mixture of subunits from both parents; or partially or fully redundant when a series of protein complexes with different orthologous members can be established because both alleles are present for some or all subunits, respectively (Timouma *et al.*, 2021). This way, biologically meaningful phenotypic predictions can be derived as a function of gene expression profiles encoding for subunits or isoforms of the involved enzymes. Regardless of the used approach to integrate omics data, the reliability of the formulated hypotheses strongly depends on the quality of gene-protein-reaction (GPR) rules included into the models, which describe how gene products concur to catalyse the associated reactions.

The *S. cerevisiae* gene IDs in the GPR rules of the iSP_1513v_0_ GSMM were replaced by the *S. pastorianus* parental alleles ID. In instances where a gene possessed both parental variants, a principle was established to acknowledge that either of the parental proteins could facilitate the reaction, given their functional redundancy. To accomplish this, an OR operator was employed to link genes that encoded different protein isoforms of the identical enzyme or subunit. The gene names were replaced with the common gene name combined with “_scer”, “_seub” or “_spast” to inform whether they are *S. cerevisiae*-like, *S. eubayanus*-like or *S. pastorianus* specific genes, respectively. This curated model, named iSP_1513, represents the *S. pastorianus* genome scale metabolic model.

The majority of the reactions are associated with genes via gene-protein-reaction (GPR) associations. In total, there are 762 reactions that are supported by only one gene, 1694 reactions that are supported by functionally redundant genes (Boolean OR), 102 reactions that are supported by a protein complex (Boolean AND) and 105 reactions that are supported by a combination of genes and protein complexes (Boolean AND and OR). However, there are also reactions that are not associated with any genes. In fact, there are 1399 reactions that are not associated with any genes of which 265 are exchange reactions. This information can be useful for understanding the genetic basis of metabolic pathways and for designing strategies for manipulating metabolic networks.

### Simulations of the metabolism of *S. pastorianus* using iSP_1513 and Yeast8

In *S. cerevisiae*, a large-scale study found that approximately 20% of the genes are essential. These genes are involved in a variety of cellular processes, including DNA replication, protein synthesis, and cell division and metabolism (Zhang & Ren, 2015). In comparison, less is known about essential genes in *S. pastorianus*, as this species has not been subjected to a large functional analysis. For the genes involved in the metabolism, we carried out single knock-out simulations using iSP_1513 and Yeast8, and predicted 34 essential and 92 essential genes, respectively. The lower number of essential genes predicted by iSP_1513 reflects the redundancy present in *S. pastorianus*, which is not captured by Yeast8. To show the importance to take into account genetic redundancy after hybridisation, we experimentally validated 2 genes (four orthologs), which are essential in *S. cerevisiae* but non-essential in *S. pastorianus*. *S. cerevisiae*-like and *S. eubayanus*-like genes of known *S. cerevisiae* essential genes, namely *FOL1* (*i.e.,* involved in folic acid synthesis) and *BPL1* (*i.e.,* encoding for a biotin protein ligase), have been deleted separately in *S. pastorianus* using long flanking PCR-mediated gene replacement. The single mutant strains were able to survive in rich medium (*i.e.,* YPD) indicating that the presence of one functional gene coming either from the *S. cerevisiae* or *S. eubayanus* parent is enough to compensate for the loss of the other (Supplementary Figure 2).

Analysis of single reaction knock-out indicated that both models have the same essential reactions with the exception for one (Table 1 and Supplementary Table 5). The virtual knock-out of hexokinase (D-glucose:ATP) r_0534 reaction (*i.e.,* promoting the transfer of a phosphate from an ATP to a D-glucose to produce D-glucose 6-phosphate) is essential in Yeast8, while is predicted to produce very low biomass using iSP_1513 (Supplementary Table 5). This is probably an artifact due to a the iSP_1513 model predicting marginally more D-fructose 6-phosphate converted into D-glucose 6-phosphate (r_0467, glucose-6-phosphate isomerase) and D-glucose 1-phophate (r_0888 phosphoglucomutase) than Yeast8, and hence feeding into the starch and sucrose metabolism. Overall, we found variation in the prediction of essential genes, but not reactions, between iSP_1513 and Yeast8.

**Table 1:** Predicted number of essential genes and reaction in SD medium + 2% glucose in *S. pastorianus* and *S. cerevisiae*.

| Model | Essentiality | SD + 2% Glucose |
| --- | --- | --- |
| iSP_1513 | Essential genes | 34 |
|  | Essential reactions | 165 |
| Yeast8 | Essential genes | 92 |
|  | Essential reactions | 166 |

We then validated our model for growth on different sugars. We predicted the growth of *S. pastorianus* using both iSP_1513 and Yeast8 in SD complete medium with different carbon sources, by performing Flux Balance Analysis with the objective function set to growth optimization. The iSP_1513 model showed biomass production in all the sugars. In glucose, maltose, maltotriose and ethanol, iSP_1513 and Yeast8 showed similar biomass produced. In medium containing melibiose as sole carbon source no biomass was detected using Yeast8, while our model was able to predict growth (Table 2, Figure 2). In fact, *S. pastorianus* has been experimentally shown to be able to utilize melibiose as a carbon source for growth and fermentation, while *S. cerevisiae* is not able to support growth on this sugar alone (Stewart, 2016), although some *S. cerevisiae* strains have been shown to metabolize melibiose to some extent (Mahilkar *et al.*, 2022). Hence, it is important to note that the growth and biomass yields of *S. pastorianus* on different sugar sources can be influenced by many factors, including the specific strain of yeast used, the composition of the culture media, and the environmental conditions of the fermentation process.

**Figure 2:**
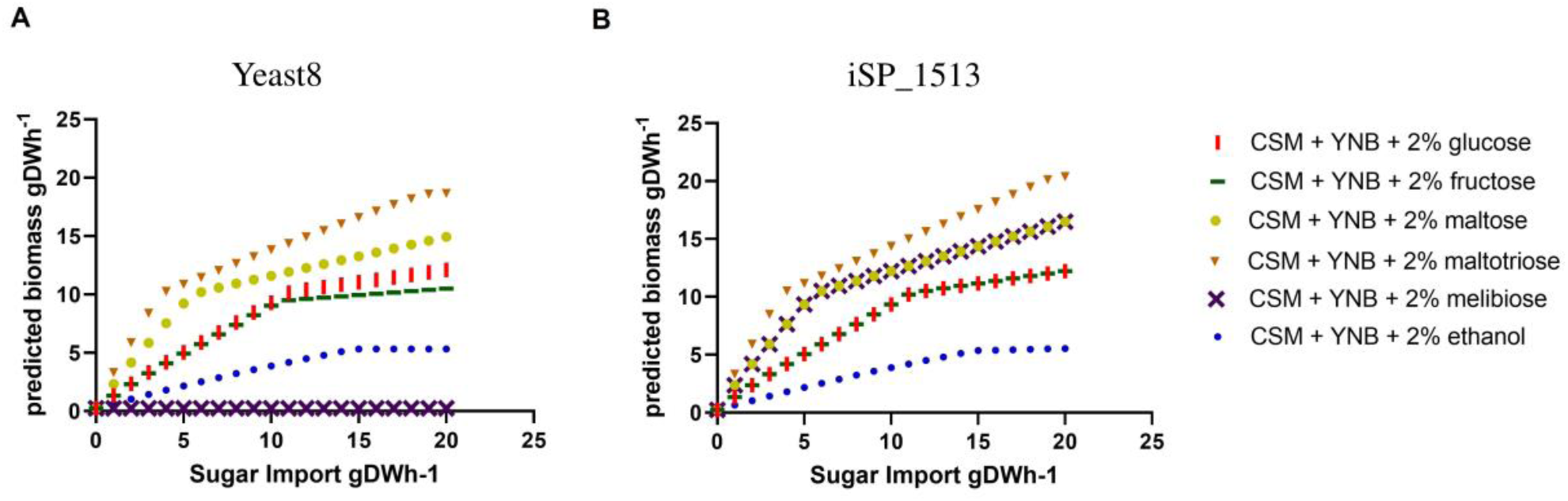
Predicted biomass (grams dry weight per hour) according to the sugar uptake. A sugar import of 20 represent the total consumption of the sugar present in the SD medium. **Panel A**: Predictions using Yeast8; **Panel B:** Predictions using iSP_1513.

**Table 2:** Predicted biomass (grams dry weight per hour) of *S. pastorianus* using iSP_1513 and Yeast8 in different media.

| Media | iSP_1513<br>(biomass gDW.h<sup>-1</sup>) | Yeast8<br>(biomass gDW.h<sup>-1</sup>) |
| --- | --- | --- |
| SD | 0.241 | 0.234 |
| SD + 2% Glucose | 2.36 | 2.35 |
| SD + 2% Maltose | 4.204 | 4.157 |
| SD + 2% Maltotriose | 5.915 | 5.859 |
| SD + 2% Melibiose | 4.204 | 0.234 |
| SD + 2% Ethanol | 1.041 | 1.034 |

### iSP_1513 metabolite secretion simulations in SD medium with various leucine content

The secretion of 2-phenyl ester (roses, honey, apple, sweet flavours), isoamyl acetate (banana and pear flavours), ethyl decanoate (floral and fruity flavours) and ethyl octanoate (apple, tropical fruit, sweet flavours) were investigated in SD medium with different concentration of leucine.

Regardless of the presence or absence of leucine, iSP_1513 model predictions indicate that *S. pastorianus* can produce 2-phenyl ester, ethyl octanoate, ethyl decanoate, isoamyl acetate, in a minimal media (Table 3). *S. pastorianus* possesses inherent metabolic capabilities that enable enhanced synthesis of these compounds. Two genes *ATF1* and *ATF2*, encoding for alcohol acetyltransferases (AATase, EC 2.3.1.84), are responsible for synthesizing acetate esters, including isoamyl acetate, through the reaction between acetyl coenzyme A and their respective alcohols (Yoshimoto *et al.*, 1999). These esters play a significant role in shaping the flavour profiles of beer and various other alcoholic beverages. *S. pastorianus* CBS 1513 possesses a *S. eubayanus*-like *ATF1* gene (SPGP0DZ02280), and both *S. cerevisiae*-like and *S. eubayanus*-like *ATF2* genes (SPGP0F01100 and SPGP0Q01180, respectively).

**Table 3:** Predictions of metabolite secretion in SD medium with various leucine content using flux variability analysis (objective set to growth, fraction of optimum 90%).

| Media | 2-phenylethyl ester | isoamyl acetate | ethyl octanoate | ethyl decanoate |
| --- | --- | --- | --- | --- |
| SD | 1.242 | 2.127 | 0.6722 | 0.6057 |
| SD without leucine | 1.271 | 1.584 | 0.6877 | 0.6197 |
| SD with extra leucine | 1.246 | 2.842 | 0.675 | 0.6082 |

The absence of leucine in the medium is predicted by the model to have an enhancing effect on the production of 2-phenyl ester, ethyl octanoate, and ethyl decanoate in *S. pastorianus*. Furthermore, higher concentrations of leucine in the medium are predicted to correspond to an increased production of isoamyl acetate. These predictions have been validated by quantifying these compounds, via GC-MS, in *S. pastorianus* cultures grown on SD, SD with an additional 100mg/L leucine and SD without leucine. When comparing the volatiles produced in the different media, we found a significantly higher amount of 2-phenyl ester, ethyl octanoate, and ethyl decanoate produced when *S. pastorianus* was grown on SD-Leu, (Figure 3A-C), and a significantly higher amount of isoamyl acetate when the medium supplemented with leucine (Figure 3D). The increase of isoamyl acetate can be attributed to leucine’s role as a precursor in the biosynthesis of isoamyl acetate via the Ehrlich pathway. The availability of leucine in the medium likely promotes the enzymatic conversion of leucine to isoamyl acetate, resulting in higher levels of isoamyl acetate production. Overall, the amount of leucine in the extracellular medium may lead to a redirection of metabolic pathways, promoting or supressing the synthesis of these volatiles. Such scenario has been correctly predicted by our model, and highlights the ability of iSP_1513 to be used in biotechnology for media engineering when growing *S. pastorianus*.

**Figure 3:**
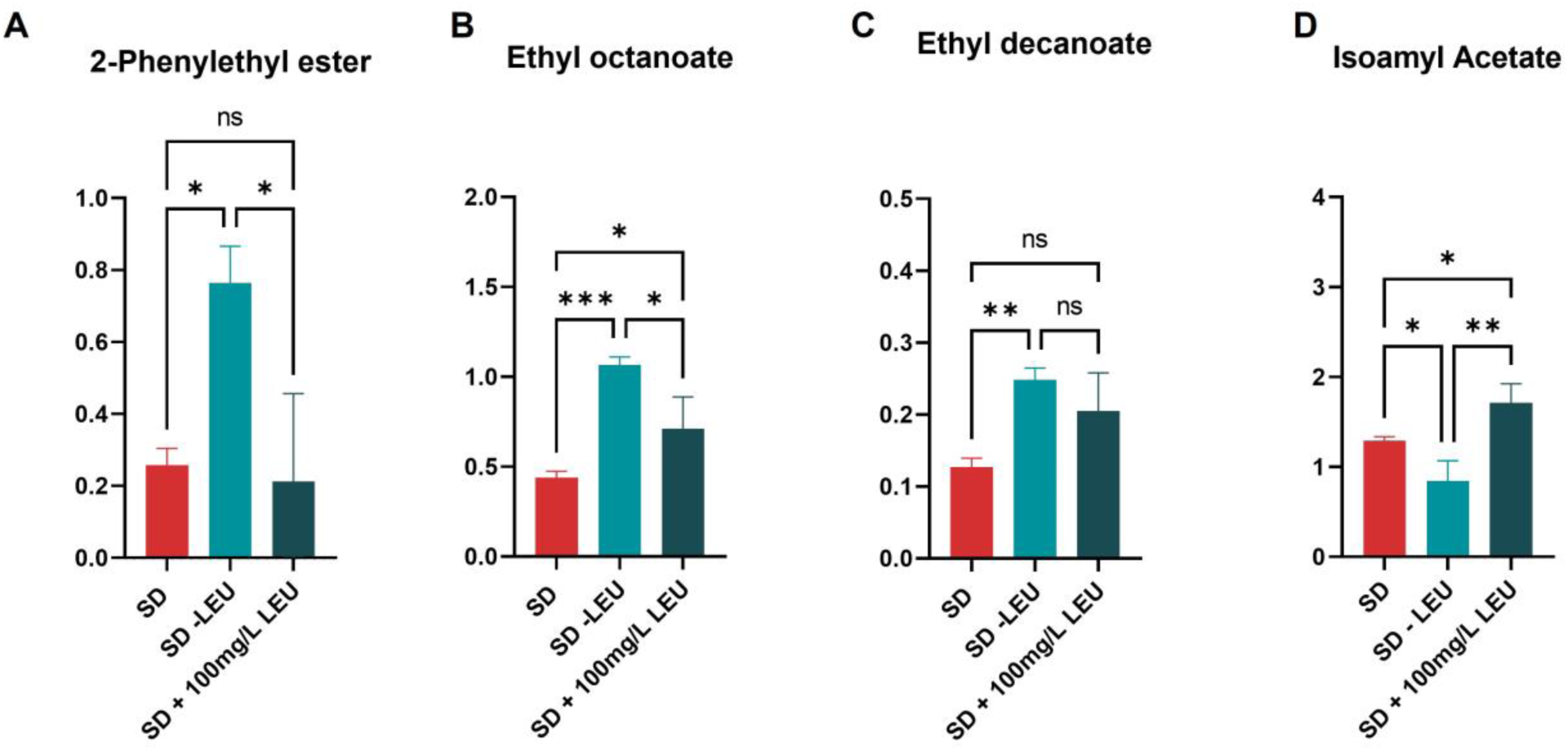
*S. pastorianus* CBS 1513 metabolite production in SD medium, SD without leucine and SD with an additional 100 mg/L leucine measured by GC-MS for the metabolites **Panel A:** ethyl octanoate; **Panel B:** ethyl decanoate; **Panel C:** 2-phenyl ester; **Panel D:** isoamyl acetate.

### Integration of transcriptome data to investigate the impact of temperature on growth

The mapping of transcriptome data to constrain genome scale metabolic model reactions allows the integration of additional parameters for the simulations such as temperature. The GSMM also provides a scaffold for the identification and visualisation of the contribution of each parental allele to metabolic pathways, uncovered by the transcription data. We have transcriptome data of *S. pastorianus* CBS1513 cultured in SD complete medium and SD medium lacking leucine at 13 °C, 22 °C and 30 °C (Timouma et al, 2021).

By mapping the normalised read counts onto the GSMM from the culture grown in SD medium at 13 °C, 22 °C and 30 °C, we found that 1107, 1139 and 1221 iSP_1513 reactions have been constrained, respectively; while for the cultures grown in SD medium lacking leucine at 13 °C, 22 °C and 30 °C, we found that 1102, 1175 and 1170 iSP_1513 reactions have been constrained, respectively.

To showcase the usefulness of this approach, we predicted using the iSP_1513 the glycerol production at 13 °C and 30 °C in SD medium, and we experimentally validated the predictions. Glycerol is important for the maintenance of membrane integrity during temperature shifts. At hot temperatures, yeast cells increase glycerol production as a protective mechanism against heat stress, while at cold temperatures, they decrease glycerol production to maintain proper membrane fluidity (Gao *et al.*, 2019). Overall, the regulation of glycerol production in yeast is dependent on the temperature, which acts as a crucial factor in maintaining membrane functionality under varying environmental conditions. In SD medium, iSP_1513 predicted that glycerol production reaches its peak at 30 °C, whereas the lowest levels are detected at 13 °C (Figure 4A). To validate these predictions, small scale fermentations were performed at 13 °C and 30 °C in SD medium. As predicted, the production of glycerol was found to be two-fold higher at 30°C compared to 13°C (Figure 4B). The utilization of transcriptome data mapping to constrain the genome-scale metabolic model reactions yielded a precise forecast of glycerol production while affirming the inclusion of temperature as a parameter in the analysis.

**Figure 4:**
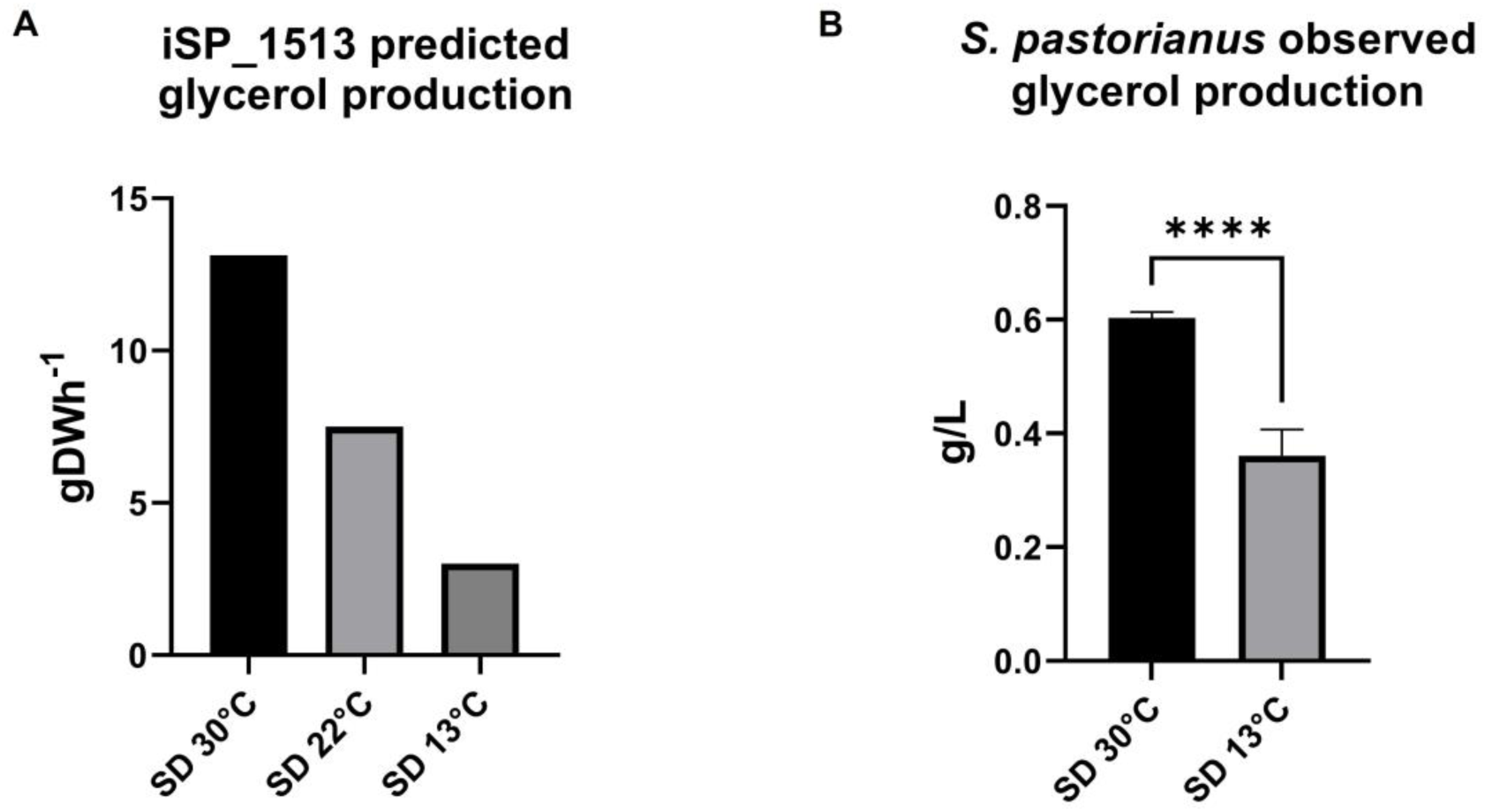
*S. pastorianus* CBS 1513 glycerol production in SD medium as predicted by the iSP_1513 model constrained using **Panel A**: the transcriptome data obtained in SD medium at 13 °C, 22 °C and 30 °C, and **Panel B**: as measured experimentally with HPLC in SD medium at 13 °C and 30 °C.

The GSMM also provides a scaffold for the identification and visualisation of the contribution of each parental allele to metabolic pathways, uncovered by the transcription data. The integration of transcriptome data to investigate the parental sub-genome activity is particularly important for hybrid species, where both parental gene copies are present and, according to the environmental conditions, only one parental gene may be expressed (Timouma *et al.*, 2021). In SD medium at 22 °C, we found that for the pentose phosphate pathway, although both parental genes can carry the reactions (Figure 5A), there is a preference of cellular function between parental alleles, with the oxidative and non-oxidative parts of pathway primarily carried out by the *S. eubayanus*-like and *S. cerevisiae*-like genes, respectively (Figure 5B, Supplementary Table 6). These results hold true also at 13 °C and 30 °C (Supplementary Table 6). Some reactions show that one of the two alleles is predominant in all the temperatures. For example, the phosphogluconate dehydrogenase reaction, supported by *GND1* or *GND2* (both having *S. cerevisiae*-like and *S. eubayanus*-like alleles), is carried out at 80%, 85% and 90% by the *S. eubayanus*-like alleles at 13 °C, 22 °C and 30 °C, respectively (Table 4). Other key metabolic reactions show a shift of parental allele usage according to the temperature. For example, the ribokinase reaction, supported by *RBK1*, is equally supported by both parental alleles at 13 °C and 22 °C, while the *S. cerevisiae*-like alleles are primarily expressed at 30 °C (Table 4). The phosphopentomutase reactions, supported by *PGM1* (*S. eubayanus*-like alleles), *PGM2* (both parental alleles) or *PGM15* (both parental alleles), is equally supported by both parental alleles at 13 °C, while at 22 °C and 30 °C there is a shift of expression toward the *S. eubayanus*-like alleles (Table 4). Finally, the phosphofructokinase reactions, supported by a protein complex composed with the product of *PFK1* (both parental allele) and *PFK2* (both parental alleles), is mainly supported by the *S. eubayanus*-like alleles at 13 °C while at 30 °C it is equally supported by both parental alleles (Table 4).

**Figure 5:**
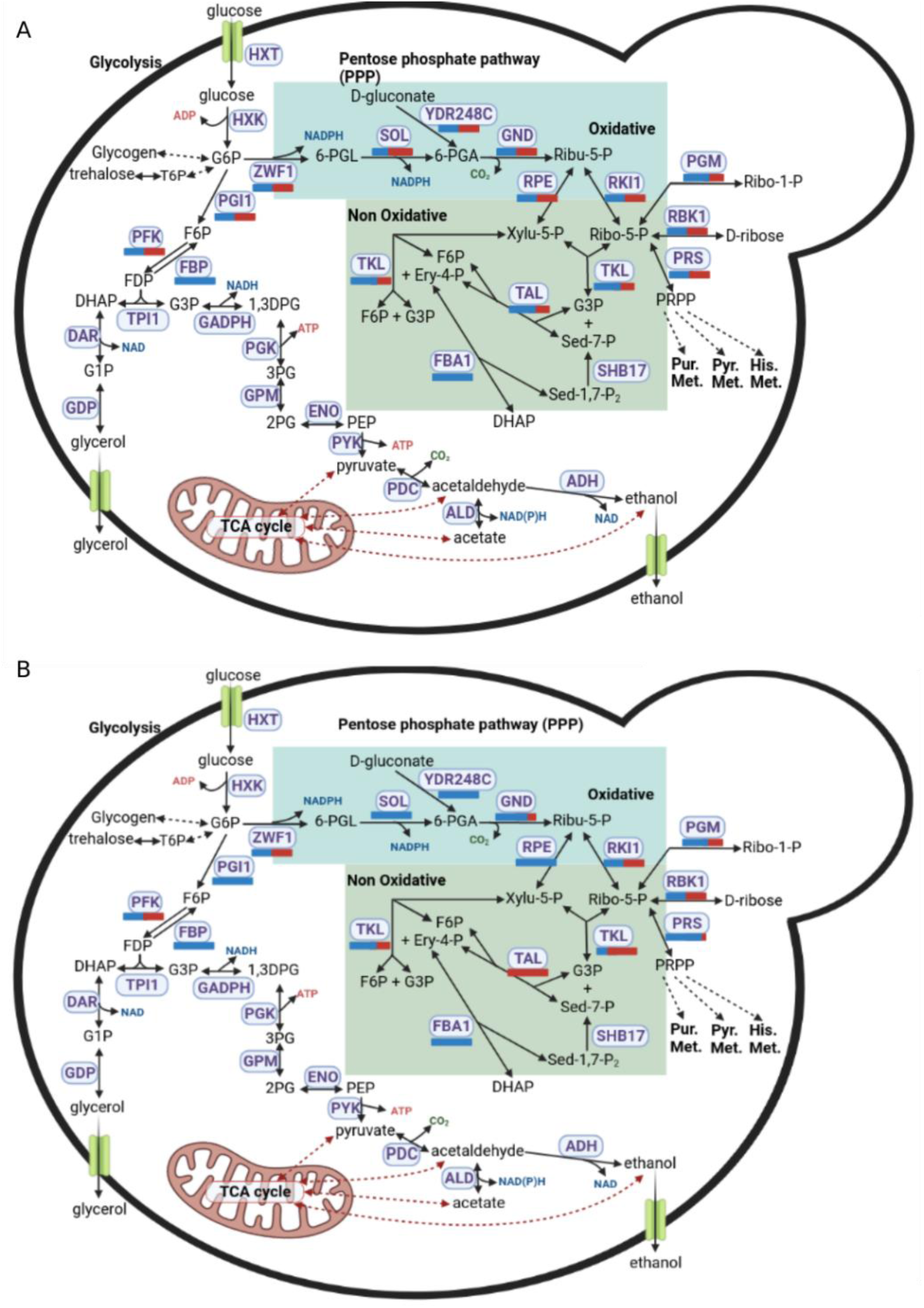
Carbon central metabolism in *S. pastorianus*. **Panel A:** Proportion of *S. eubayanus*-like and S. *cerevisiae*-like genes present in the genome supporting each reaction. **Panel B:** Proportion of *S. eubayanus*-like and S. *cerevisiae*-like genes expressed at 22 in SD medium. *S. eubayanus*-like and S. *cerevisiae*-like genes are represented as blue and red bars, respectively.

**Table 4:** Proportion of *S. eubayanus*-like and *S. cerevisiae*-like genes supporting reactions of the pentose phosphate pathway in SD medium at 13 °C, 22 °C and 30°C.

| Reaction name | GSM reaction GPR | Proportion of genes supporting the reaction in SD media at 13°C |  | Proportion of genes supporting the reaction in SD media at 22°C |  | Proportion of genes supporting the reaction in SD media at 30°C |  |
| --- | --- | --- | --- | --- | --- | --- | --- |
|  |  | *S. cerevisiae*-like genes | *S. eubayanus*-like genes | *S. cerevisiae*-like genes | *S. eubayanus*-like genes | *S. cerevisiae*-like genes | *S. eubayanus*-like genes |
| glucose-6-phosphate isomerase | PGI1_Scer or PGI1_Seub | 0% | 100% | 0% | 100% | 0% | 100% |
| glucose-6-phosphate dehydrogenase | ZWF1_Scer or ZWF1_Seub | inconclusive | inconclusive | inconclusive | inconclusive | inconclusive | inconclusive |
| 6-phosphogluconolactonase | SOL1_Scer or SOL1_Seub or SOL2_Scer or SOL3_Scer or SOL3_Seub or SOL4_Scer or SOL4_Seub or | 0% | 100% | 0% | 100% | 0% | 100% |
| phosphogluconate dehydrogenase | GND1_Scer or GND1_Seub or GND2_Scer or GND2_Seub | 20% | 80% | 15% | 85% | 10% | 90% |
| ribulose 5-phosphate 3-epimerase | RPE1_Scer or RPE1_Seub | 0% | 100% | 0% | 100% | 0% | 100% |
| transketolase | TKL1_Seub or TKL2_Scer or TKL2_Seub | 85% | 15% | 85% | 15% | 85% | 15% |
| transaldolase | TAL1_Seub or NQM1_Scer or NQM1_Seub |  | 100, 0 |  | 100, 0 |  | 100, 0 |
| ribose-5-phosphate isomerase | RKI1_Scer or RKI1_Seub | inconclusive | inconclusive | inconclusive | inconclusive | inconclusive | inconclusive |
| ribokinase | RBK1_Scer or RBK1_Seub | 50% | 50% | 50% | 50% | 55% | 45% |
| phosphoglucomutase | PGM1_Seub or PGM2_Scer or PGM2_Seub | inconclusive | inconclusive | inconclusive | inconclusive | inconclusive | inconclusive |
| phosphopentomutase | PGM1_Seub or PGM2_Scer or PGM2_Seub or PRM15_Scer or PRM15_Seub | 50% | 50% | 40% | 60% | 30% | 70% |
| phosphoribosylpyrophosphate synthetase | (PRS1_Seub and (PRS2_Scer or PRS2_Seub)) or (PRS1_Seub and (PRS3_Scer or PRS3_Seub)) or (PRS1_Seub and (PRS4_Scer or PRS4_Seub)) or ((PRS2_Scer or PRS2_Seub) and (PRS5_Scer or PRS5_Seub)) or ((PRS4_Scer or PRS4_Seub) and (PRS5_Scer or PRS5_Seub)) | 5% | 95% | 5% | 95% | 5% | 95% |
| ATP:D-Gluconate 6-phosphotransferase | YDR248C_Scer or YDR248C_Seub | 0% | 100% | 0% | 100% | 0% | 100% |
| Fructose-bisphosphate aldolase | FBA1_Seub | inconclusive | inconclusive | inconclusive | inconclusive | inconclusive | inconclusive |
| Fructose-bisphosphatase | FBP1_Seub | 0% | 100% | 0% | 100% | 0% | 100% |
| phosphofructokinase | ((PFK1_Scer or PFK1_Seub) and (PFK2_Scer or PFK2_Seub)) | 40% | 60% | 45% | 55% | 50% | 50% |

## CONCLUSIONS

The development of iSP_1513 represents a significant advancement in modelling the metabolism of hybrid yeast strains and enhance industrial approaches to fermentation processes. Despite the challenges posed by the lack of molecular data and the complexity of physiological features of yeast aneuploid hybrids, recent progress in genome sequencing and annotation has made possible the construction of genome-scale metabolic models for specific *S. pastorianus* strains.

iSP_1513, the first GSMM tailored to the industrial strain *S. pastorianus* CBS 1513, addresses the issue of functional redundancy caused by orthologous parental alleles. This unique characteristic enables accurate predictions of gene knock-outs and omics data mapping. Notably, contrarily to the Yeast8 model, iSP_1513 is able to utilize melibiose as a carbon source. Moreover, iSP_1513 precited that *S. pastorianus* produces a higher amount of 2-phenyl ester, ethyl octanoate, and ethyl decanoate flavour compounds when grown on SD medium lacking leucine, and a significantly higher amount of isoamyl acetate when the medium was supplemented with leucine.

We developed a universal Python3 function as a library, crucial for mapping transcriptome data onto GSMM reactions, hence facilitating growth predictions under various environmental conditions including temperature as parameter. iSP_1513 correctly predicted that the production of glycerol increases when the external temperature increases (*i.e.,* accordance with the experimental validation). In SD medium, we found that there is a preference of cellular function between parental alleles, with the oxidative and non-oxidative parts of the pentose phosphate pathway primarily carried out by the *S. eubayanus*-like and *S. cerevisiae*-like genes, respectively. For some reactions of this pathway, a shift of parental allele usage according to the temperature was observed.

Overall, iSP_1513 represents a significant step forward in our ability to model the metabolism of hybrid yeast strains and optimize industrial fermentation processes. The iSP_1513 model provides a platform for predicting optimal growth conditions in different environments, thanks to the inclusion of the functional redundancy of parental alleles in enzymatic reactions within the Gene Protein rules of the model. By employing Flux Balance Analysis (FBA) and Flux Variability Analysis (FVA), iSP_1513 can predict growth in diverse environments and is capable of anticipating reactions that promote or suppress the production of specific aroma compounds. This feature enables more efficient and tailored fermentation processes, making it a valuable tool in the field of biotechnology and industrial bioprocessing. Ultimately, the iSP_1513 model can be used in combination with evolutionary algorithms to predict adaptation trajectories under different environmental pressures, and hence provide a top-down framework to study genome evolution in hybrids.

## MATERIALS AND METHODS

### Genome sequence and annotation

*S. pastorianus* CBS 1513 strain has been sequenced and assembled (Hewitt *et al.*, 2014; Okuno *et al.*, 2016). Its genome sequence is available from the National Center for Biotechnology Information (NCBI: txid1073566). The Yeast Genome Annotation Pipeline (YGAP) has been used to predict the potential Open Reading Frames (ORFs) in its genome (Proux-Wéra *et al.*, 2012). HybridMine tool (https://github.com/Sookie-S/HybridMine, v4.0) has been used to identify the parental allele content in this strain (Timouma *et al.*, 2020). Among the 9,728 potential ORFs of *S. pastorianus* CBS 1513, there are 5,218 *S. eubayanus*-like alleles and 3,739 *S. cerevisiae*-like alleles, of which 1706 are ortholog alleles (Timouma *et al.*, 2020). To enrich the pool of *S. pastorianus* gene annotated, one-to-one orthologs were searched between *S. pastorianus* predicted proteins and the proteome of the two parental species, using HybridMineP (https://github.com/Sookie-S/HybridMineP). HybridMineP is using the same pipeline as HybridMine except for using BLASTP instead of BLASTN for finding one-to-one orthologs between the species.

### GSMM draft reconstruction

The COBRApy package have been used in a Python 3.6 environment to reconstruct the genome scale metabolic model. The yeast consensus genome-scale model Yeast8 version 8.6.2 (Lu *et al.*, 2019), has been used as a template to construct a draft genome scale model of *S. pastorianus* CBS1513. Yeast8 contains 1160 genes, 2744 metabolites and 4063 reactions. First, the *S. cerevisiae*-like alleles of *S. pastorianus* were mapped. Next, one-to-one orthologs were searched between the *S. eubayanus*-like alleles of *S. pastorianus* and the *S. cerevisiae* S288C genes using HybridMine software (Timouma *et al.*, 2020). The gene-protein-reaction (GPR) associations, which are Boolean statements connecting genes to reactions, were updated to take into account the ortholog parental alleles found in *S. pastorianus*.

Additionally, to identify *S. pastorianus* specific reactions, a draft GSMM reconstruction were generated through the KBase (Arkin *et al.*, 2018) narrative interface. *S. pastorianus* CBS1513 (NCBI: txid1073566) genome in FASTA format was uploaded into the Staging Area and, subsequently, imported into the narrative through the ‘Import FASTA File as Assembly From Staging Area’ app. Its genome annotation file (Timouma *et al.*, 2020) in GFF3 format was uploaded into the Staging Area and, subsequently, imported into the narrative through the ‘Import GFF3/FASTA File as Genome From Staging Area’ app. Draft metabolic reconstructions were generated through the ‘Build Fungal Model’ app and exported in SBML format through the ‘Bulk Download Modeling Objects’ app. KBase draft GSMM doesn’t take into account the gene redundancy, the redundant orthologous alleles were added to the reaction GPR rule using an in-python script. The *S. pastorianus* genes and reactions that are present in the KBase GSMM draft but absent from iSP_1513 were identified. Genes in the KBase model absent in the iSP_1513 draft were searched as they could be potentially added. The reactions supported by these genes were mapped to the iSP_1513 GSMM to identify genes/reactions specific to *S. pastorianus*. With that methodology, genes/reactions were added to iSP_1513. Gprofiler tool (Reimand *et al.*, 2016) was used to conduct a gene ontology enrichment analysis on the removed genes.

The *S. pastorianus* model has been named iSP_1513 and is encoded in the Systems Biology Markup Language (SBML).

### Model simulations

A metabolic flux corresponds to the amount of a metabolite processed by one or more catalytic steps per unit of time, normalized by cellular abundance. Metabolic fluxes have a unit of gram dry weight (gDW) (Stephanopoulos *et al.*, 1998). Simulation using stoichiometric models allow a quantitative understanding of metabolism. Flux balance analysis (FBA) calculates the flow of metabolites through a metabolic network, therefore the growth rate of an organism or the production rate of a metabolite can be predicted, under the assumption that during exponential growth, the metabolic function produces a constant flux of biomass (Orth *et al.*, 2010). Steady-state mass balance is assumed. FBA is concerned with the following linear program (LP):

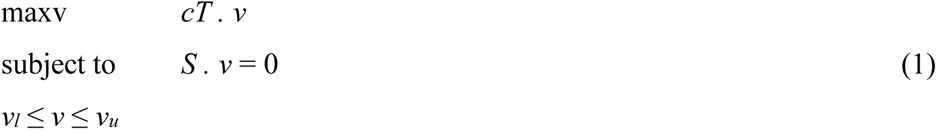

where *S* is and *m * n* stoichiometric matrix of a metabolic network with *m* metabolites and *n* reactions, and *c* is the vector representing the linear objective function; *v* is the rate of metabolic flux, with *V* ⊆ ℝ^n^; the vectors *v_l_* and *v_u_* represent the flux of the lower and upper bounds, respectively. The upper bounds were set as 1000 while the lower bounds were set as -1000 for the reversible reactions and 0 for the irreversible reactions. The substrate uptake reaction (such as the consumption of glucose, oxygen, or ammonia) is changed to a specific value according to the medium of cell growth. The constraints *S . v* = 0 together with the upper and lower bounds specify the feasible region of the problem.

While FBA only finds the maximum flux for the model reactions, flux variability analysis (FVA) (Mahadevan & Schilling, 2003) calculates the minimum and maximum flux for the reactions while maintaining some state of the network, e.g., supporting 90% of maximal possible biomass production rate. FVA allows the exploration of alternative optima of (1). Let *w* represent some biological objective such as the biomass production. After solving (1) with *c* = *w*, FVA solves two optimization problems for each flux *v_i_* of interest

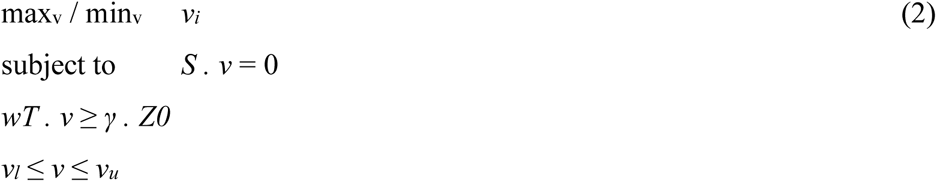

where *Z0* = *wT*.*v0* is an optimal solution to (1), *γ* is a parameter, which controls whether the analysis is done *w.r.t*. suboptimal network states (0 ≤ *γ* < 1) or to the optimal state (*γ* = 1). Assuming that all n reactions are of interest, FVA requires the solution of 2n LPs.

Flux balance analysis (FBA) and Flux variability analysis (FVA) simulations were performed using COBRApy package (v. 0.18.1). The IBM ILOG CPLEX solver has been used to find the optimal and sub-optimal solutions. FBA and FVA were performed on conditions reflecting synthetic defined (SD) medium + 2% glucose (Supplementary Table 7). SD medium composition was taken from Harrison *et al.*, 2007 and the substrate intake fluxes were updated to model the SD medium purchased from Formedium that was used experimentally.

### Transcriptome data processing

Gene expression data of *S. pastorianus* CBS1513 grown in SD medium + 2% glucose and SD medium without leucine +2% glucose at 13 °C, 22 °C and 30 °C were taken from Timouma *et al.*, 2021. The raw read counts of the RNAseq data were normalised for sequencing depth and gene length using the Transcripts Per Kilobase Million (TPM) method, using the bioinfokit python 3 library. The read counts were divided by the length of each gene in kilobases, which gives reads per kilobase (RPK) normalisation. All the RPK values in a library were counted up and the result was by 1,000,000, which gives the “per million” scaling factor. Finally, the RPK values were divided by the “per million” scaling factor, which gives the TPM. The advantage of that method is that the sum of all TPMs in each library are the same and the genes within a library can be compared.

### Mapping transcriptome data to constrain GSMM reactions

To integrate temperature as a parameter in addition to the media composition, gene expression data of *S. pastorianus*, normalised with the TPM method, were mapped to the metabolic reactions following the methodology described in Lee *et al.*, 2012. The Boolean OR relationship signals alternative catalysts so the total capacity of the reaction is given by the sum of its components, while the Boolean AND relationship signals a complex between several gene products therefore the maximum complex concentration is given by the minimum concentration of its components. The AND Boolean relationship is above the OR relationship within the GPR. Irreversible reactions will have their upper bound constrained with the positive final expression value while reversible reactions will have both their upper bound and lower bound constrained with the positive and negative final expression values, respectively.

We developed a universal python 3 function, “map_transcriptome_data”, that can be broadly used for any genome scale model. First, “map_transcriptome_data” analyses the GPR associations in a model of metabolic reactions. The function categorizes the GPR associations into several groups based on their Boolean relationships, such as OR, AND, and a mixture of both. The reactions for which the GPR that is supported by only one gene will be constrained using the expression data of the gene. The GPRs that contains only the OR Boolean relationship will be constrained by the sum of the expression data of the genes. The GPRs that contains only the AND Boolean relationship will be constrained by the minimal expression data of the genes. The GPR that contains a mixture of OR and AND Boolean relationships are the most complex case therefore a recursive function was implement within the “map_transcriptome_data” function to tackle these cases by order of priority. A detailed explanation is presented in Supplementary File 2. Constraining reactions based on gene expression levels is a common practice in genome-scale metabolic modelling. In this case, reactions have been constrained only if all the genes within the GPR have an expression value above 10. This means that the expression level of all genes associated with a given reaction must be greater than 10 in order to constrain the reaction. The threshold of 10 is chosen as it is assumed to be above the noise level of gene expression data, which can be influenced by technical and biological variability.

“map_transcriptome_data()” requires 4 positional arguments: ’model’, ’transcriptomeData’, ’threshold_abundance’, and ’max_bound’, with ’model’ being the genome scale model, ’transcriptomeData’ a dictionary containing the gene IDs (as written in the genome scale model) as keys and the transcription levels as values, ’threshold_abundance’ the threshold to consider that a gene is expressed rather than be noise (for example 10 as recommended in DESeq2 documentation), ’max_bound’ the value of the upper bounds when there is no restriction (for example in the Yeast8 model the lower and upper bounds range from -1000 to 1000 so the ’max_bound’ is 1000). This function can be installed using python3 pip or downloaded at https://github.com/Sookie-S/Mapping-of-transcriptome-data-to-genome-scale-scale-model-reactions. A description of how to install and use this function is provided in Supplementary File 2.

### Media and yeast culture

*S. pastorianus* CBS1513 and *S. cerevisiae* BY4743 strains were used in this study. Starter cultures were generated by inoculating 5ml volume of YPD (yeast extract, 10 g/L, peptone, 20 g/L, glucose, 20 g/L) with cells and incubating at 22 °C with shaking at 200rpm overnight. Biomass was recorded via optical density measurement at 600 nm with an BioSpectrometer (Eppendorf).

### PCR–based generation of gene knockout mutants

For gene deletions in *S. pastorianus*, a PCR -based fragment fusion method approach was used (Oakley *et al.*, 2007; Zhao *et al.*, 2019). Gene knockout cassettes were created by amplification of and Phusion of three fragments, 1Kb homolog region upstream and downstream of the target gene and the selectable marker. Primers used for amplification are listed in Supplementary Table 8. Strains were transformed using the LiAc/SS carrier DNA/PEG method (Gietz & Schiestl, 2007) and transformants were confirmed by analytical PCR.

### HPLC analysis

*S. pastorianus* was grown in 200mL in Synthetic Minimal Media (SD medium: 1X Yeast Nitrogen Base-YNB; 1X Complete Supplement Mixture-CSM, both Formedium; 2% (w/v) glucose) after inoculation with a washed overnight culture to reach an initial OD of 0.1. Cells grew for 7 and 10 days at 30°C and 13°C, respectively prior collection to identify the metabolites at the end of the fermentation. The concentration of metabolites, such as glucose and glycerol were measured by HPLC using a 1260 Infinity II LC System with a Refractive Index Detector (Agilent). A 300 × 7.8 mm Hi-Plex HPLC Column (Agilent) was equilibrated with 5 mM H_2_SO_4_ in HPLC grade water at 55° at a 0.8 mL/min flow rate. Prior to analysis, the samples were filtered (pore size 0.45 μm). Quantification was achieved using a RID-detector. Calibration curves from authentic standards were used to quantify metabolites produced.

### GC-MS analysis

Volatile compounds were detected and analysed by GC-MS. Cells were grown in 5 mL of SD complete, SD lacking leucine (SD-LEU) and SD with 100mg/L Leucine on a 20 mL vials added with 25 μL of the internal standard 2-octanol (2.5 mg/L) after inoculation with a washed overnight culture to reach an initial OD of 0.1. All the cultures grew at 20°C for 10 days.

Samples were analysed on an Agilent 7890B Gas Chromatograph (GC) paired with an Agilent 5977B Series Mass Selective Detector (MSD) and operated with a Gerstel MPS dual head system. Vials containing the sample were incubated (30 °C, 10 min) to preconcentrate volatile analytes into the headspace. Followed by extraction for 5 minutes with a 100 um PDMS fibre and injection into the front inlet. The GC separation was performed using an Agilent VF-5MS column (30m x 25mm x 0.25uM) with a flow rate of 1mL·min^-1^. The oven gradient was: 40 °C held for 2 min, 15°C·min^−1^ to 300°C and held for 5 min. The total run time for each sample was 24.3 minutes. The inlet was set at 280 °C, the transfer line was kept at 300°C, the EI source at 230 °C and the quadrupole at 150 °C. The MSD mass range scanned was *m/z* 40-400.

The acquired data was processed using LECO ChromaTOF. Peak picking, peak annotation, and statistic confirmation were performed, and peak identification was completed by checking the linear temperature programmed retention index (LTPRI), which is available in the NIST RI database. Inter-measurements peak alignment was performed based on the retention times and mass spectrum. An inter-class comparison was performed between sample class and blank class to eliminate artifact compounds.

## ACKNOWLEDGMENTS

The authors wish to thank Katherine Hollywood and Maira Hernandez-Guzman for their assistance with the GC-MS and HPLC experiments, respectively. The authors thank Neil Swainston and Doug Kell for their initial guidance on the genome scale metabolic model reconstruction and simulations.

## AUTHOR CONTRIBUTIONS

ST developed the genome scale metabolic model, the python3 function for mapping the transcriptome data, and carried out the gene deletion experiments. LNBC performed the HPLC and GC-MS experiments. All the author analysed and discussed the data. DD and JMS supervised the research. All authors wrote the manuscript, have read and agreed to the published version of the manuscript.

## DATA AVAILABILITY

iSP_1513 genome scale metabolic model is available on GitHub at https://github.com/Sookie-S/Genome-scale-metabolic-model-of-S.-pastorianus-CBS-1513-iSP_1513-.

## CODE AVAILABILITY

The python3 library developed to map transcriptome data into a genome-scale metabolic model reactions, to enable the imposition of restrictions that accurately represent the experimental conditions being investigated, is available at https://github.com/Sookie-S/Mapping-of-transcriptome-data-to-genome-scale-scale-model-reactions.

## SUPPORTING FIGURES LEGEND

**Supplementary Figure 1:** GO term enrichment (molecular function) of the 56 *S. cerevisiae* specific genes that were removed from Yeast8 model.

**Supplementary Figure 2:** Growth in YPD agar of *S. pastorianus* CBS 1513 wild type (WT), and *S. pastorianus* heterozygote mutant strains carrying the deletion of *S. cerevisiae*-like *FOL1* (“S. cer like *FOL1*△”), *S. eubayanus*-like *FOL1* (“S. eub like *FOL1*△”), *S. cerevisiae*-like *BPL1* (“S. cer like *BPL1*△”) and *S. eubayanus*-like *BPL1* (“S. eub like *BPL1*△”).

