## Supplementary figures and images for "Development of a genome scale metabolic model for the lager hybrid yeast *S. pastorianus* to understand evolution of metabolic pathways in industrial settings"

### Supplementary Fig.1

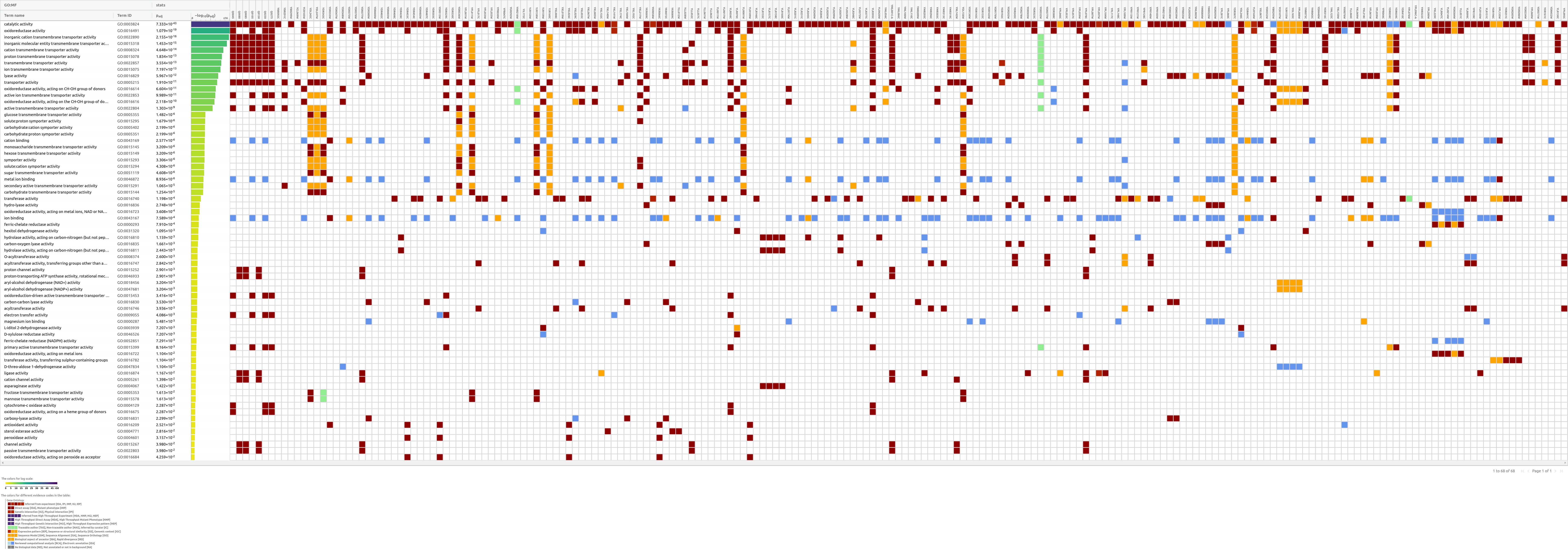

### Supplementary Fig.2

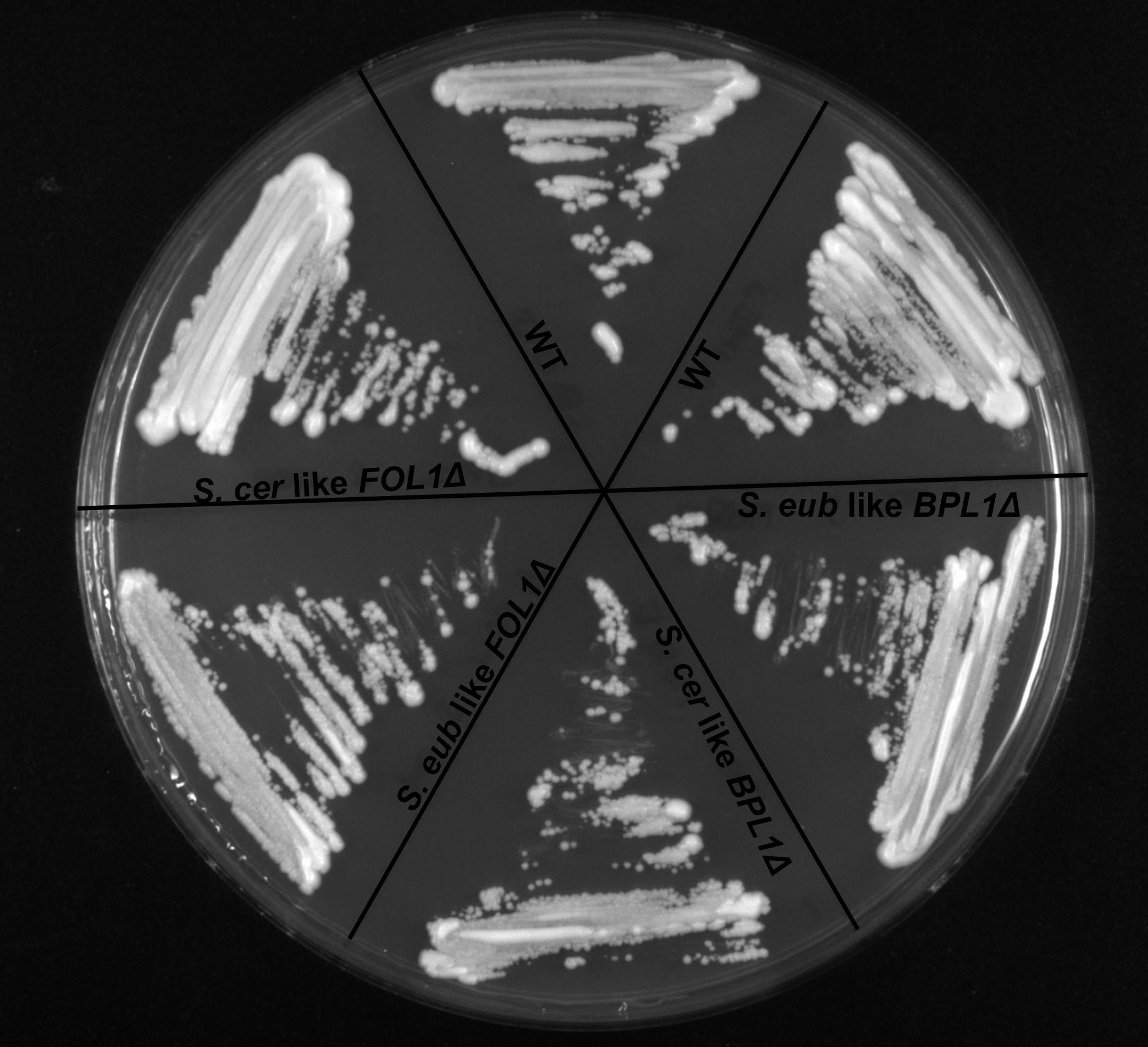
