## Supplementary File 1 for "Development of a genome scale metabolic model for the lager hybrid yeast *S. pastorianus* to understand evolution of metabolic pathways in industrial settings"

**Supplementary File S1**

**Genes, reactions and metabolites added to the iSP_1513 model**

The cytosolic reaction r_4712 AMP nucleosidase reaction, supported by a protein complex composed by three subunits, has been added to iSP_1513.

H_2_O (c) + AMP (c) --> Adenine (c) + alpha-D-Ribose 5-phosphate (c) (r_4712)

All the genes supporting this reaction are already present in the model except for the *S. eubayanus*-like gene SPGP0R01370, which has been added.

Three nitrilase reactions, r_4713, r_4714 and r_4715, supported by a protein complex composed by three subunits (*NIT1*, *NIT2* and *NIT3*) were added to iSP_1513.

2.0 H_2_O (c) + Indole-3-acetonitrile (c) --> Indoleacetate (c) + NH_4_^+^ (c) (r_4713)

2.0 H_2_O (c) + 2-Aminopropanenitrile (c) --> L-Alanine (c) + NH_4_^+^ (c) (r_4714)

2.0 H_2_O (c) + 4-Amino-4-cyanobutanoic acid (c) --> L-Glutamate (c) + NH_4_^+^ (c) (r_4715)

The metabolites Indole-3-acetonitrile, 2-Aminopropanenitrile and 4-Amino-4-cyanobutanoic were added to the model as they are metabolites of the reactions r_4713, r_4714 and r_4715, respectively. The *S. cerevisiae*-like gene SPGP0DD00150 (NIT1_Scer) was also added to the model, while *NIT2* and *NIT3* are already present in the model.

The reaction acetate reversible transport via proton symport r_4716 has been added alongside the *S. cerevisiae*-like gene SPGP0AE00950 (*BPH1*_Scer) that supports it.

Acetate (e) + H^+^ (e) <=> Acetate (c) + H^+^ (c) (r_4716)

The GPR of the dolichyl-phosphate-mannose--protein mannosyltransferase reaction r_0362 was updated. The *S. cerevisiae*-like gene SPGP0F00900 (*PMT6*) was added to it forms a dimer with SPGP0G00180 (*PMT4*).

The GPR of the glutamine-fructose-6-phosphate transaminase reaction r_0477 was updated. The *S. cerevisiae*-like gene SPGP0DB02080 and the *S. eubayanus*-like gene SPGP0S02100 were added to the GPR of the reaction.

The GPR of the 1,3-beta-glucan synthase reaction r_0005 was updated was updated with the *S. cerevisiae*-like genes SPGP0DB00800 (*GAS3*_Scer), SPGP0K00380 (GAS4_Scer), SPGP0K01360 (GAS5_Scer) and the *S. eubayanus*-like genes SPGP0T00270 (GAS1_Seub), SPGP0N00740 (GAS2_Seub), SPGP0P02990 (GAS4_Seub) and SPGP0P02040 (GAS5_Seub).

The GPR of the asparagine synthase (glutamine-hydrolysing) reaction r_0211 was updated with the *S. eubayanus*-like gene SPGP0DX00470_Seub.

The GPR of the endopolygalacturonase reaction r_ 0365 was updated with the *S. eubayanus*-like gene SPGP0R03420_Seub.

The GPR of the D-mannose transport, D-fructose transport, D-glucose transport and D-galactose transport reactions r_ 1139, r_1134, r_1166, r_1135 respectively were updated with the *S. pastorianus*-specific genes SPGP0B01550_Spas and SPGP0R03480_Spas.

The GPR of the maltose transport reaction r_1227 was updated with the *S. eubayanus-*like and *S. pastorianus*-like genes SPGP0DN00100_Seub and SPGP0P03210_Spas.

The GPR of the homocysteine S-methyltransferase reaction r_0544 was updated with the *S. pastorianus*-like gene SPGP0U02250_Spas.

The GPR of the 6-phosphogluconolactonase reaction r_0091 was updated with the *S.cerevisiae*-like genes SPGP0M03420 (SOL1_Scer) and SPGP0AE01280 (SOL2_Scer), and *S. eubayanus*-like gene SPGP0DV00410 (SOL1_Seub).

The GPR of the phosphoglycerate mutase reaction r_0893 was updated with the *S. cerevisiae*-like genes SPGP0C02080 (GMP2_Scer) and SPGP0K01120 (GMP3_Scer).

The GPR of the V-ATPase Golgi and V-ATPase vacuole reaction r_1085 and r_1086 respectively were updated with the *S. pastorianus*-specific gene SPGP0P01340 (VMA10_Spas)

The GPR of the chitin deacetylase reaction r_0271 was updated with the *S. eubayanus*-like gene SPGP0N01030.
