## Supplementary File 2 for "Development of a genome scale metabolic model for the lager hybrid yeast *S. pastorianus* to understand evolution of metabolic pathways in industrial settings"

**Supplementary File S2.**

**Description of the algorithm to map transcriptome data.**

The function "map_transcriptome_data" classifies the associations described in the Gene-Protein-Reaction (GPR) rules based on their Boolean relationships, namely OR, AND, and a combination of both. In cases where a reaction is supported by a GPR associated with only one gene, the expression data of that gene is utilized to impose constraints. GPRs that exclusively involve the OR Boolean relationship are constrained by the summation of expression data from the associated genes. Conversely, GPRs featuring solely the AND Boolean relationship are constrained by the minimum expression data among the associated genes. The most complex scenario arises when a GPR involves a combination of OR and AND Boolean relationships. To address such cases in a prioritized manner, a recursive function has been implemented within the "map_transcriptome_data" function.

For example, as illustrated in the following hypothetic GPR (1):

*GENE_ASSOCIATION: ((A or B) and C) or (D and E)) and F*	(1)

with *A, B, C, D, E* and *F* different genes, that as genes expression values 100, 70, 30, 60, 90 and 50 respectively.

The recursive function searches for the first block that is not nested within the GPR (1), starting from the end of the GPR, and will calculate the final expression value according to the Boolean operator. In the example, it corresponds to the block (*D* and *E*):

*GENE_ASSOCIATION: ((A or B) and C) or (D and E)) and F*	(2)

As the Boolean AND relationship is between D and E, the expression value corresponds to the minimum of their expression values:

*(D and E): min(D, E) = min(60,90) = 60*	(3)

The block (D and E) in (2) is next replaced by 60 (3). Recursively, the block that is the least nested becomes (*A* or *B*):

*GENE_ASSOCIATION: ((A or B) and C) or 60) and F*	(4)

As the Boolean OR relationship is between A and B, the expression value corresponds to the sum of their expression values:

*(A or B): (A + B) = (100 + 70) = 170*	(5)

The block (A or B) in (4) is next replaced by 170 (5). Recursively, the block that is the least nested becomes (170 and C):

*GENE_ASSOCIATION: ((170 and C) or 60) and F*	(6)

As the Boolean AND relationship is between 170 and C, the expression value corresponds to the minimum of their expression values:

*(170 and C): min(D, E) = min(170, 30) = 30*	(7)

The block (170 and C) in (6) is next replaced by 30 (7). Recursively, the block that is the least nested becomes (30 or 60):

*GENE_ASSOCIATION: (30 or 60) and F*	(8)

As the Boolean OR relationship is between 30 and 60, the expression value corresponds to the sum of their expression values:

*(30 or 60): (30 + 60) = 90*	(9)

The block (30 + 60) in (8) is next replaced by 90 (9). Recursively, the block that is the least nested becomes (90 and F):

*GENE_ASSOCIATION: 90 and F*	(10)

As the Boolean AND relationship is between 90 and F, the expression value corresponds to the minimum of their expression values:

*90 and F: min(90, F) = min(90, 50) = 50*	(11)

The block (90 and F) in (10) is next replaced by 50 (11).

*GENE_ASSOCIATION: 50*	(12)

The recursive function stops when no more blocks are detected (12).

**Installation instructions for the algorithm to map transcriptome data**

The "map_transcriptome_data" function can be installed using python3 pip as following:

‘python3 -m pip install GSMM_transcriptome_data_mapper’

This function can be used with the following command lines on a python3 terminal:

‘from GSMM_transcriptome_data_mapper import transcriptome_mapper’

‘transcriptome_mapper.map_transcriptome_data(model,transcriptomeData,threshold_abundance,max_bound)’

The script can also be downloaded at <https://github.com/Sookie-S/Mapping-of-transcriptome-data-to-genome-scale-scale-model-reactions>.
